# Dissociable Roles for Working Memory in Sensorimotor Learning

**DOI:** 10.1101/290189

**Authors:** Samuel D. McDougle, Jordan A. Taylor

## Abstract

The computations underlying cognitive strategies in sensorimotor learning are poorly understood. Here we investigate such strategies in a common sensorimotor transformation task. We show that strategies assume two forms, which are thought to reflect distinct working memory representations: discrete caching of stimulus-response contingencies (e.g., a look-up table), and time-consuming parametric computations (e.g., mental rotation). Subjects’ reaction times and errors suggest that both strategies are employed during learning, and trade off based on the amount of practice and the complexity of the task. Experiments using pressured preparation time support these dissociable working memory representations: In discrete response caching, time pressure elicits multi-modal distributions of movements; in parametric mental rotation, time pressure elicits a shifting distribution of movements between visual targets and distal goals, consistent with analog re-computing of a movement plan. Furthermore, a generalization experiment reveals that discrete and parametric strategies produce, respectively, more localized or more global transfer effects. These results describe how qualitatively distinct working memory representations are leveraged for motor learning and have downstream consequences for behavioral flexibility.

## INTRODUCTION

When first learning a new motor skill, the process of selecting an appropriate action often involves a time-consuming, deliberative process. Consider someone first learning to play the piano: Ideally, she could quickly learn a stimulus-response mapping relating notes on the staff to their appropriate keys. However, as anyone who has attempted to learn a new instrument knows, learning this mapping is tractable when a musical score has a few notes in a small range, but as it gets more complicated, things fall apart — if we just consider just the number of keys in an octave, it easily exceeds normal working memory capacity (Miller, 1956). A common strategy (used in piano pedagogy) to overcome this limitation is to approach it parametrically: She can anchor her thumb on middle C and reference other notes on the lines of the musical staff relative to this key. While this strategy affords her the ability to play a more complex melody within a few minutes of practice, it becomes increasingly cumbersome the further a given note is from middle C — echoing the phenomena of mental scanning and mental rotation (Kosslyn et al., 1978; Shepard and Metzler, 1971). These two strategies, one a discrete map (caching stimulus-response pairs) and one a flexible algorithm (computing relative distances from C), offer two ways to approach learning a novel motor skill.

In simpler motor tasks, like the visuomotor rotation task (Krakauer, 2009), subjects often leverage strategies to rapidly improve performance (Taylor et al., 2014). These strategic processes appear to be related to higher reaction times (Haith et al., 2015a), improved task performance (Fernandez-Ruiz et al., 2011), and the direction of eye gaze (Rand and Rentsh, 2015; de Brouwer et al., 2018). Increased reaction time, and the fact that strategies are often verbalizable, suggests that they reflect deliberative, controlled processing rather than automatic processing (Cohen et al., 1990). Control processes often rely on working memory, making it one candidate system that may underlie cognitive strategies for motor learning. Indeed, evidence suggests that spatial working memory ability correlates with performance in visuomotor tasks (Seidler et al., 2012), and both recruit similar neural circuits (Anguera et al., 2010). Moreover, spatial working memory ability correlates with the use of explicit strategies in visuomotor rotation tasks (Christou et al., 2016). However, it remains unclear what kinds of working memory representations are used for strategies in motor learning.

Here we set out to directly characterize how working memory may manifest in a visuomotor rotation task, which requires subjects to adapt to sensory feedback that is rotated relative to their direction of movement. We hypothesized that deliberate working memory strategies would take two broad forms, either discrete response caching (RC) or parametric mental rotation (MR). A discrete RC strategy is here defined as the maintenance of acquired associations between a set of stimuli and a set of responses maintained in memory (Collins and Frank, 2012; Provost et al., 2013), perhaps relying on processing in prefrontal cortex (Petrides 1985; Curtis & D’Esposito, 2003). As a form of item-based working memory, the efficacy of a discrete RC strategy should be subject to load (e.g. the number of items to be stored; Cowan, 2010; Collins and Frank, 2012).

A parametric MR strategy, on the other hand, is the canonical example of an “analog” computation (Shepard & Metzler, 1971; Cooper, 1976), in which an internal mental representation is manipulated in visual working memory in a manner similar to an external physical object (Hyun & Luck, 2007). Evidence from behavior and neurophysiology provides support for mental rotation as a computation in the planning of reaching movements: Reaction time (RT) during the mental rotation of a reaching direction scales with the magnitude of required rotation, sharing remarkable similarities to the mental rotation of visual percepts (Georgopoulos & Massey, 1987; Pellizzer & Georgopoulos, 1993; Bhat & Sanes, 1998). Moreover, decoded neuronal population vectors in motor cortex appear to “rotate” through directional space during mental rotation of a reach plan (Georgopoulos et al., 1989; but see Cisek & Scott, 1999). Critically, mental rotation can compute rotations with arbitrary signs or magnitudes, and through multiple planes (Shepard & Metzler, 1971). Mental rotation thus constitutes a flexible algorithm that is formally equivalent to the application of a rotation matrix to some mental representation. Similar algorithms likely exist for other sensorimotor transformations. For instance, linear RT effects are observed when humans compute gain perturbations on reaching extent (Bhat & Sanes, 1998); here, the heuristic would simply represent a scalar transformation rather than a rotation, though the same logic applies. The idea of mental rotation specifically describing explicit learning in visuomotor rotation tasks has been suggested before (Fernandez-Ruiz et al., 2011), but here we provide the first direct test of this idea.

An additional question concerns how different representations change over the course of motor learning. Recent work on classic visual mental rotation supports the idea of parametric versus discrete strategies in that domain - if subjects are exposed to many unique objects in a visual mental rotation task, RT signatures of mental rotation persist over days; however, if they are only exposed to a few images during extended training, mental rotation effects diminish with time, suggesting a shift to an item-based retrieval strategy (Provost et al., 2013). Similarly, recent work on visuomotor rotation learning suggests that time-consuming strategic learning processes appear to become more automatic with practice (Huberdeau et al., 2017). Together, these findings are broadly consistent with Logan’s (1988) theory of skill automatization, where learning proceeds from an algorithmic stage to an associative memory-retrieval stage, the latter requiring repeated practice of specific “instances.” Our experiments here are poised to confirm this transition as a model of the cognitive aspects of visuomotor learning, directly test the putative computations underlying these different stages, and also test their downstream consequences. In *Experiment 1*, we present evidence in support of distinct working memory representations for motor learning, and provide empirical support for a transition from algorithmic to retrieval-based strategies in motor learning. In *Experiments 2* and *3*, we expose the putative within-trial mechanisms underlying these distinct strategies. In *Experiment 4*, we characterize downstream consequences of different learning representations on behavioral flexibility and generalization.

## RESULTS

### Experiment 1: Dissociable working memory strategies in a visuomotor adaptation task

Subjects performed a standard visuomotor rotation task (see Methods; for a review see Krakauer, 2009), where visual feedback was rotated relative to the reaching direction of the hand (Fig. 1A). Our hypothesis was that in low set size conditions (small number of targets), subjects would primarily use a discrete response caching (RC) strategy (i.e., a look-up table), while in high set size conditions they would use a parametric mental rotation (MR) strategy (i.e., a rule-based algorithm). We grouped subjects via a 2 × 2 between-subjects factorial design, crossing the factors Set Size and Rotation Magnitude. For Set Size, subjects were exposed to either 2 or 12 possible target locations during the task: In the 2T condition, one of two targets, separated by 180°, was presented on each trial. The precise location of targets were randomized and counterbalanced across subjects. In the 12T condition, target locations were pseudorandomly presented at one of 12 possible locations. For Rotation Magnitude, subjects either experienced a 25° or 75° rotation during the rotation block. Varying rotation magnitude was necessary to test for the operation of a parametric MR strategy. Critically, feedback, in the form of a small visual cursor, was only provided at the end of the reach and was delayed by 2 s, a manipulation which has been shown to limit implicit motor adaptation (Kitazawa et al., 1995; Brudner et al., 2016; Schween et al., 2017; Parvin et al., 2018) and thus better isolate strategic learning.

**Figure 1:**
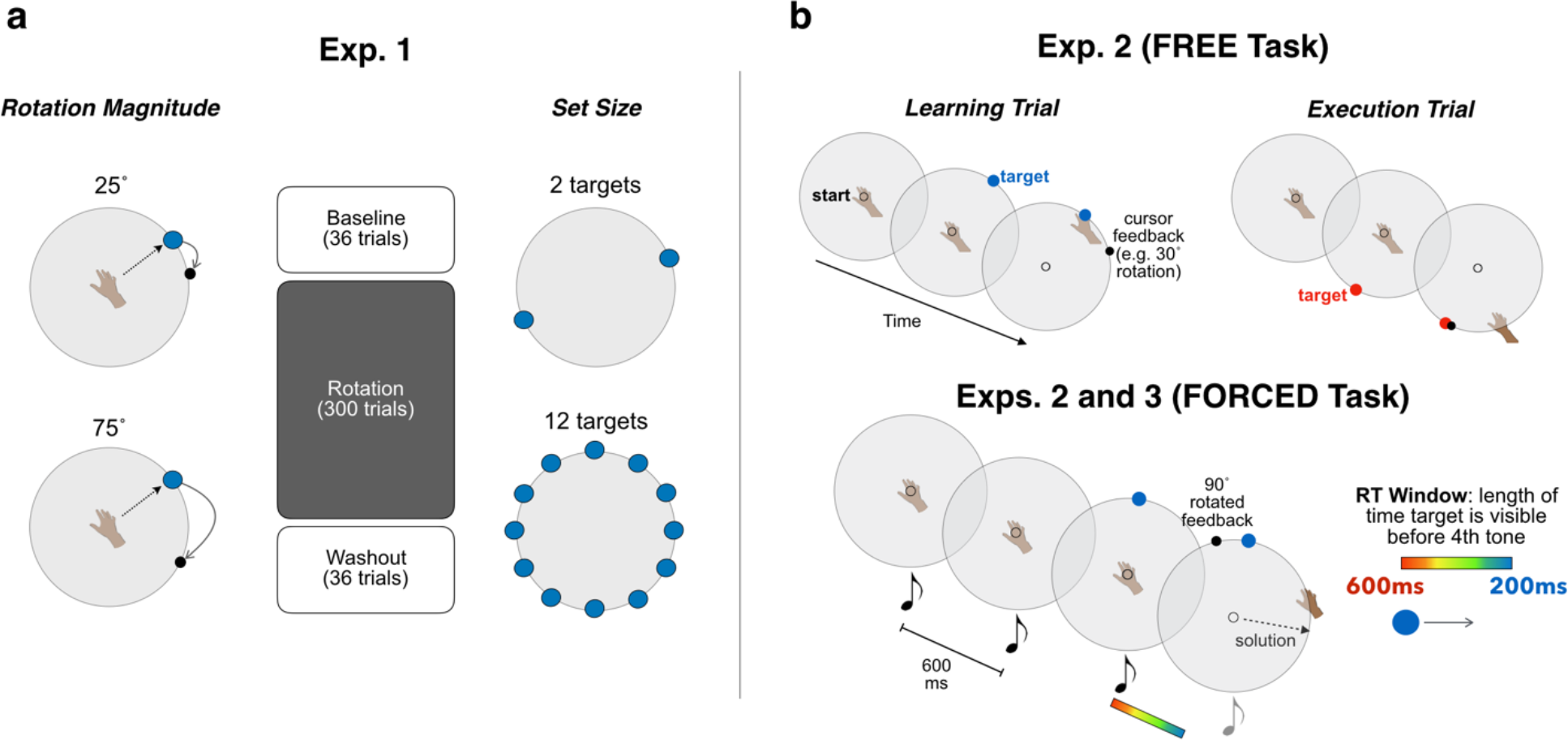
Experiment 1-3 task design. (**A**) *Experiment 1:* Subjects performed a standard visuomotor rotation task, attempting to land a rotated visual cursor on a target. A between-subjects, 2 X 2 design was used, crossing the factors Rotation Magnitude (the size of the rotation in the rotation block; left) and Set Size (the number of possible target locations in the task; right). (**B**) *Experiments 2-3:* FREE task: Subjects performed trial pairs consisting of learning trials (left) and execution trials (right). Thirteen rotation sizes were pseudorandomly presented (−90°:15°:90°). FORCED task: Subjects were required to initiate their movement after target presentation and < 100 ms after the fourth tone. Targets could appear in either one of 12 locations (Exp. 2) or one of 2 locations (Exp. 3). The time of target appearance was titrated to induce subjects to react with a distribution of RTs.

Our first analysis focused on reaction time (RT), a standard metric for quantifying the operation of executive processes (Cohen et al., 1990). We hypothesized that in the high Set Size group (12T), RT would be modulated by Rotation Magnitude throughout learning, consistent with the idea that subjects were parametrically mentally rotating their motor plans on each trial. However, in the low Set Size group (2T), we expected this effect only early in learning, while subjects discovered the “structure” of the task. Thus, we expected RTs to converge across rotation magnitudes in the 2T groups due to the caching of discrete responses. Our 2 X 2 design allowed us to test these specific predictions: parametric MR in all groups early in learning, and a shift to RC for the two 2T group later in learning.

Consistent with these predictions, we observed that rotation magnitude affected RT in all groups early in learning, but only the 12T groups late in learning (Fig. 2A). To quantify this, we examined the means of median RTs over 6 cycles between early learning (first 6 cycles) and late learning (last 6 cycles) across our four conditions. Note, we defined a cycle as 2 trials in the 2T group and 12 trials in the 12T group to control for the inherent difference between set size conditions in the number of exposures at each target location. These RTs (Fig. 2C) were submitted to a 3-way mixed factorial ANOVA with a within-subjects factor of Time (early vs. late learning), and between-subject factors of Set Size (2T vs. 12T) and Rotation Magnitude (25° vs. 75°). We note that the lack of Set Size effects early in learning are an example of how an algorithmic strategy can essentially break Hick’s Law, which describes a log-linear increase in RT due to the number of stimuli subjects are prepared to react to (Hick, 1952).

**Figure 2:**
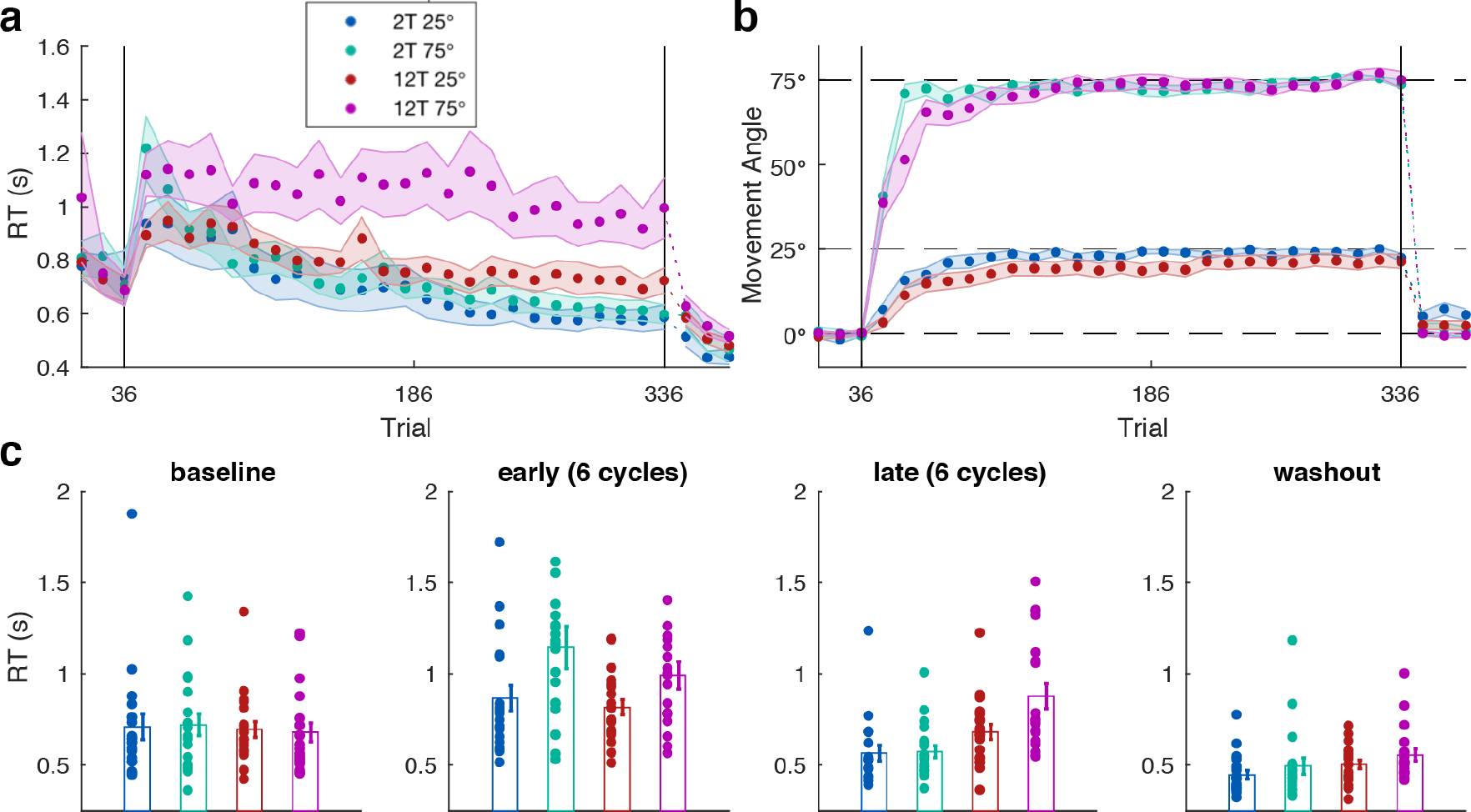
Experiment 1 results. (**A**) Time course of mean RTs, averaged over 2-trial and 12-trial cycles in the 2T and 12T groups. Vertical lines delineate the task block (baseline, rotation, and washout). (**B**) Movement angle across the groups, averaged over 12-trial bins. (**C**) Mean of median RTs averaged over the cycles of trials in the beginning (early) and end (late) of learning, and the full baseline and washout blocks (see Methods). Error bars = 1 s.e.m.

We found a significant main effect of Time (*F*(76) = 67.97, *p* < 0.001), reflecting the fact that all groups decreased their RT over training. A significant effect of Rotation Magnitude was also observed, suggesting that subjects may have employed a parametric MR strategy — larger rotations resulted in larger RTs (*F*(76) = 8.32, *p* = 0.005). We did not find a main effect of Set Size (*F*(76) = 0.90, *p* = 0.34), but critically found a significant two-way Time X Set Size interaction (*F*(76) = 20.87, *p* < 0.001) and a three-way Time X Set Size X Rotation interaction (*F*(76) = 4.56, *p* = 0.036). *Post hoc t*-tests (Bonferroni-corrected) revealed an effect of Rotation Magnitude on late learning RT only in the 12T groups (*t*(38) = 2.45; *p* = 0.019), but not in the 2T groups (*t*(38) = 0.15; *p* = 0.88). Taken together, these results are consistent with the hypothesis that all groups may have used a parametric MR strategy to find the task solution early in learning, and then either maintained that strategy for the remainder of the task (12T group), or transitioned to a discrete RC strategy, where RT was no longer modulated by rotation size (2T group). These results are consistent with a recent study on practice and set size effects in classic visual mental rotation (Provost et al., 2013), suggesting that general-purpose spatial cognition strategies may support learning various types of skills.

We note that the mean of median RTs over the baseline revealed no significant difference with respect to Set Size or Rotation Magnitude (*p* = 0.63 and *p* = 0.97, respectively). While we did not expect any differences as a function of Rotation Magnitude — as the rotation had yet to be imposed — there could have been baseline differences due to Set Size as subjects experienced fewer trials at each target location in the 2T groups, and may have had to perform an additional visual search to locate the changing target location in the 12T groups. Likewise, in the washout block we found no main effect of Rotation Magnitude on RT (*F*(76) = 2.40; *p* = 0.13) and only a marginal effect of Set Size (*F*(76) = 3.35; *p* = 0.07). Given this trend in the washout block (Fig. 2C), one speculation is that differences in RT at the end of training carried over into the washout, perhaps due to habitual rather than task-related factors (Wong et al., 2017).

The observed differences in RT could not be attributed to differences in task performance between conditions: All groups displayed similar asymptotic performance, with some subtle differences in the rate of learning (Fig. 2B). To quantify this, we computed the average movement angular error (i.e., rotation - movement angle) over the first 6 cycles (early learning) and last 6 cycles (late learning) for all groups and submitted these values to a two-way repeated measures ANOVA. We found a significant main effect of Time on movement error (*F*(76) = 74.70, *p* < 0.001), reflecting learning. We found no significant main effects of Rotation (*F*(76) = 0.23, *p* = 0.64) or Set Size (*F*(76) = 1.25, *p* = 0.27), nor an interaction (*F*(76) = 1.34, *p* = 0.25). We also found no interaction between Rotation X Time (*F*(76) = 1.18, *p* = 0.28). We did, however, find a significant Time X Set Size interaction (*F*(76) = 6.28, *p* < 0.014), likely reflecting the slight learning advantage seen in the 2T groups. Lastly, there were no significant differences in movement error over the baseline block with respect to Set Size or Rotation Magnitude (*p* = 0.96 and *p* = 0.66, respectively). Taken together, the general lack of significant results on movement error suggests that neither rotation magnitude nor set size significantly affected task performance, consistent with previous results (Bond and Taylor, 2015; McDougle et al., 2017).

We now consider the role of implicit motor adaptation in our task: First, delayed feedback appeared to be effective for limiting implicit adaptation, as average aftereffects across the sample were only 2.71 ° (Fig. 2B). This suggests that the vast majority of learning was driven by a strategy rather than implicit motor adaptation (Taylor et al., 2014). However, even though aftereffects were subtle, there were significant differences in movement angle in the washout block with respect to both Set Size and Rotation Magnitude (*F*(76) = 16.10, *p* = 0.002 and *F*(76) = 9.95, *p* < 0.001, respectively), though there was no significant interaction (*F*(76) = 1.67; *p* = 0.20). Corrected t- tests revealed that only the 2T 25° group showed statistically reliable aftereffects (*t*(19) = 3.49; *p* 0.001; all other *p’s* > 0.09). Parsimony would suggest this effect was not due to implicit adaptation, as in that case significant aftereffects should also be present in the other conditions (Bond and Taylor, 2015). This result is, however, consistent with “use-dependent” learning, which describes a bias toward repeated movement directions: Use-dependent learning would be more robust in the 2T setting because only two responses are repeated, and could thus create a stronger bias (Verstynen and Sabes, 2011).

#### Distributions of reach errors reveal strategy differences

A secondary movement analysis also comports with dissociable discrete and parametric working memory strategies. We analyzed subjects’ “sign” errors (i.e. trials where subjects reached in the wrong clockwise/counter-clockwise direction relative to the correct response). Normatively, discrete RC and parametric MR strategies make disparate predictions about sign errors: In discrete RC, sign errors should represent trials where participants accidentally extract the wrong response held in working memory (i.e. a working memory “swap” error; Bays, 2016). This would constitute a −155° error in the 25° condition, and a −105° error in the 75° condition. In parametric MR, however, sign errors would most likely represent trials where participants accidentally “flip” the rotation angle, thus aiming with the correct magnitude relative to the target but in the incorrect direction (Christou et al., 2016). This would constitute a −25° error in the 25° condition, and a −75° error in the 75° condition.

As sign errors were relatively rare (especially in the 2T groups), we pooled sign error data over all subjects in each group. Sign errors were designated as trials where the subject reached ≤−15° from the target location (i.e. in the direction opposite the correct positive response). As depicted in the histograms in Figure 3, swap errors were the more common sign error in the 2T groups, consistent with an incorrect discrete response being retrieved from working memory. Flip errors, however, were the more common error in the 12T groups, consistent with a parametric algorithm that can occasionally confuse the rotation sign. (We note that given the sparsity of these data and the necessity of pooling, this is a qualitative analysis that should be interpreted with appropriate caution.)

**Figure 3:**
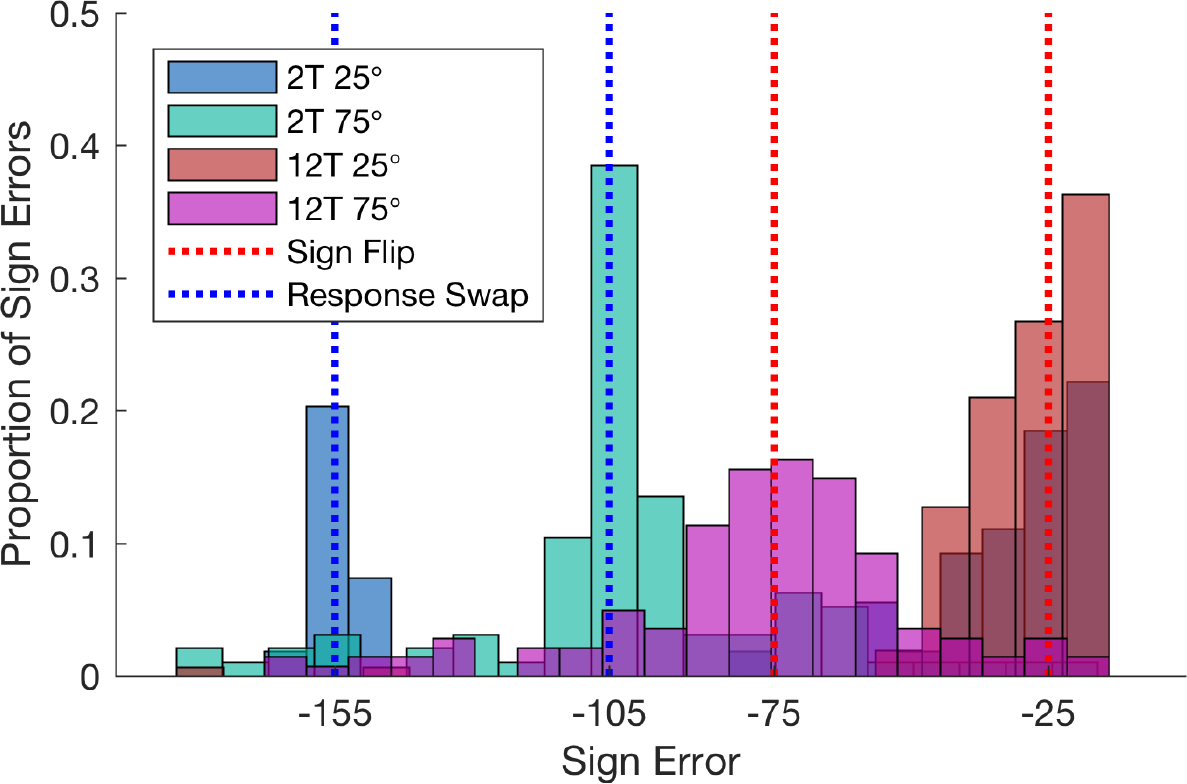
Sign error analysis. Histograms of pooled “sign error” data in each group. Vertical dotted lines represent predicted locations of “swap” errors (generating the wrong response given two choices) and “flip” errors (reaching relative to the correct magnitude of rotation but the wrong sign).

### Experiment 2: Constraining RT reveals parametric transformation of movement plans

*Experiment 1* provided evidence supporting our hypothesis that parametric MR and discrete RC strategies comprise two deliberate action selection strategies in a visuomotor adaptation task. In *Experiments 2* and *3*, we wanted to further confirm that these mechanisms were valid descriptions of true within-trial mental computations. To do this, we adopted a “forced reaction time” task (Ghez et al., 1997), which limits the amount of time subjects have to prepare their responses. This procedure has recently been used to show how different learning processes proceed during motor learning (Haith et al., 2015a; Huberdeau et al., 2017). Here, however, we wanted to use rapidly-induced movements to “read out” cognitive processes within a trial. Our specific hypotheses were as follows: Interrupting a parametric MR computation should induce intermediate movements that represent partially-rotated movement plans (*Experiment 2*), but interrupting a discrete RC computation should induce bimodal movement distributions with modes at each cached movement direction (*Experiment 3*).

To investigate mental rotation-like computations, we used a within-subject design with two tasks: One task (the FREE task) had no constraints on RT — this task was designed to replicate previous findings that, during putative mental rotation of a reaching movement, RTs are a linear function of rotation magnitude (Georgopoulos & Massey, 1987; Pellizzer & Georgopoulos, 1993; Bhat & Sanes, 1998). The second task (the FORCED task) used a forced-RT procedure to reveal the within-trial dynamics of a parametric MR strategy.

In the FREE task, subjects performed a series of trial pairs (Fig. 1B, top): In the first trial of each pair, the “learning” trial, subjects were instructed to reach directly at the displayed target and observe where the cursor landed. In the second trial of the pair, the “execution” trial, subjects were told to apply what they learned about the relationship between their movement and the resultant feedback and attempt to make the cursor terminate within the target (target positions were different between the learning and execution trials; see Methods). Rotations applied in the learning trials ranged from −90° to 90° by 15° intervals and were pseudo-randomly presented throughout the task. We hypothesized that RT would be a linear function of subjects’ reaching angles as they counteracted rotations of varying magnitude (Georgopoulos & Massey, 1987; Bhat & Sanes, 1998).

In the FREE task, subjects were generally accurate on execution trials, with reach angles on the execution trials tracking the magnitude of the rotation imposed on the corresponding learning trial (Fig. 4A; *t*-test on regression slopes: *t*(31) = 30.57, *p* < 0.001). As predicted, RT on execution trials was linearly scaled by the angle of movement produced relative to the target (Fig. 4B; *t*-test on regression slopes: *t*(31) = 6.59, *p* < 0.001), consistent with a mental rotation (MR) process. These results replicate findings that cognitive re-planning of a reaching vector displays the psychophysical standard applied to the mental rotation of visual percepts (Georgopoulos & Massey, 1987; Bhat & Sanes, 1998).

**Figure 4.**
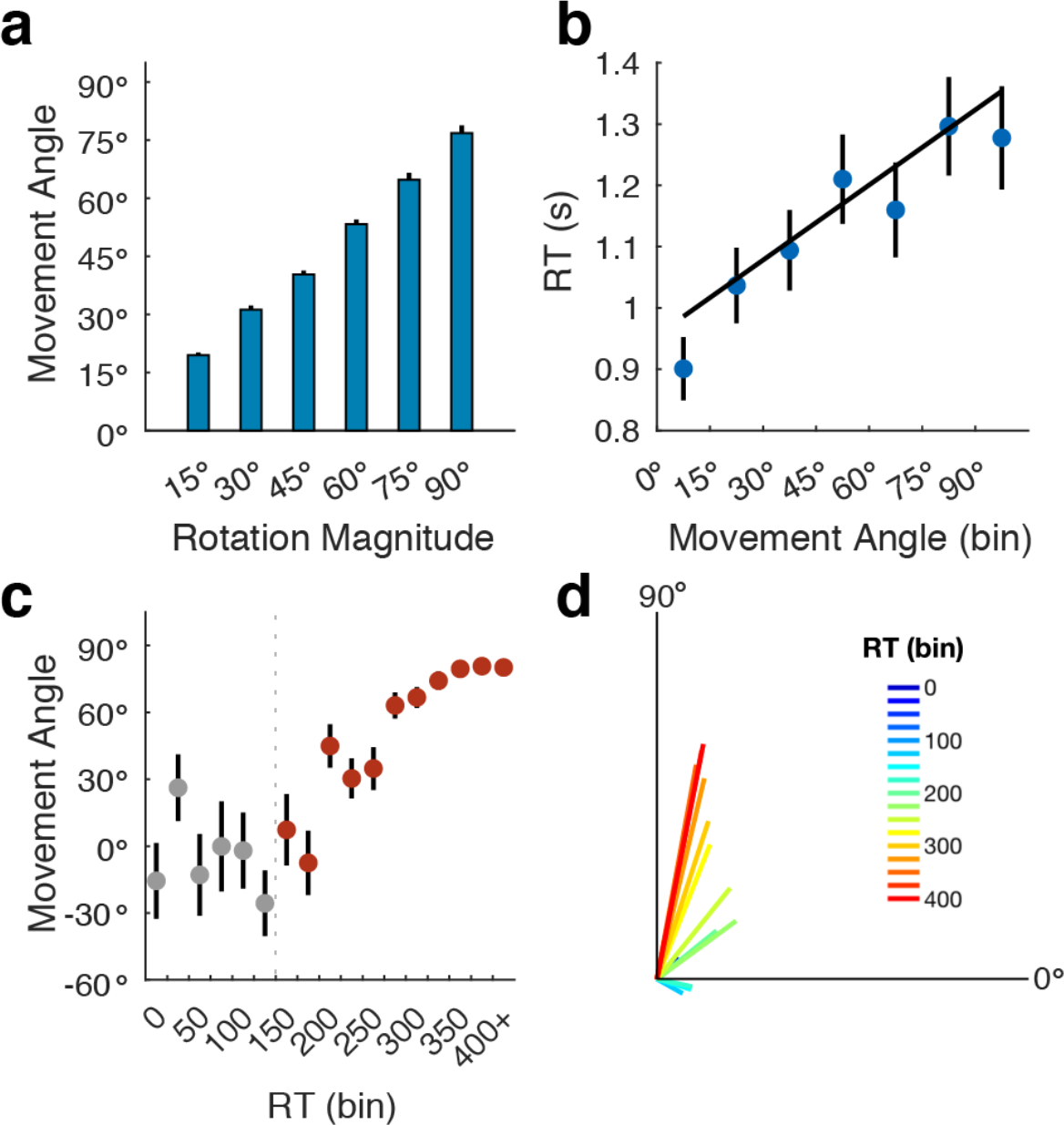
Experiment 2 results. (**A**) Absolute movement angles as a function of absolute rotation magnitude in the FREE task. (**B**) Mean of median RTs as a function of movement angle (binned by 15°). (**C**) Circular mean of movement angles as a function of RT (binned in 25 ms bins) in the FORCED task. Gray dots represent RT bins where movements were deemed as primarily random (see Methods). (**D**) Pooled vector representation of data in (**C**). Error bars = 1 s.e.m.

The FORCED task utilized a modified forced-RT paradigm to interrupt putative mental rotation (Ghez et al., 1997; Haith et al., 2016; Fig. 1B, bottom), with the prediction that movement angle would be a linear function of RT. (Thus, between the FREE and FORCED tasks, we effectively switched the independent and dependent variables.) On each trial, a countdown of four audio tones was played, and subjects were instructed to try and synchronize the initiation of their reach with the fourth tone (see Fig. 1B, bottom, and Methods). Targets could appear in one of 12 locations. The moment of target appearance was titrated such that subjects had varying amounts of time with which to compute the target location, plan, and execute their movements. For the rotation trials, a fixed rotation of 90° (or −90°, for counterbalancing) was imposed on the cursor, and subjects were thoroughly educated about the rotation before this block began.

We focused on trials where subjects reached on time in accordance with the fourth tone (*μ* = 78.90% of trials). Our first analysis involved measuring subjects’ reach angles as a function of RT. RTs were binned by 25 ms from 0 ms through 400 ms, with the final bin including all RTs above 400 ms. To quantify MR, we first needed to identify the RT at which subjects’ movement angles “settled” on a direction that was reliably sensitive to the presented target location (at very short RTs, movements should be directed randomly since there is insufficient time to process the target; Haith et al., 2016). To determine this “critical” RT, we looked at variance in movement angles in each successive 25ms RT bin. Circular variance first significantly decreased (*t*(31) = 2.28, *p* = 0.02; Supplementary Fig. 1) from the 7th to the 8th RT bin (150-175ms), suggesting that at RTs over 150ms, subjects began to make non-random movements. We note that this result is consistent with previous work (Haith et al., 2016).

After this critical bin, reaching angles linearly increased with RT towards the solution (Fig. 4C; *t*(31) = 14.17, *p* < 0.001). Figures 4C and 4D depict the results as, respectively, circular means, and the circular means and vector lengths of the pooled data. This result echoes the observed rotation of a population vector in the motor cortex, and strongly suggests a mental rotation operation during the RT (Georgopoulos et al., 1989). Moreover, the observed linear trend of intermediate movement directions (Fig. 4C, D) likely represents a behavioral correlate of intermediate states of mental rotation, confirming a fundamental assumption of parametric computations (Shepard & Metzler, 1971; Cooper, 1976).

#### Mental rotation paces are correlated between FREE and FORCED tasks

Our conclusions assume that the FREE and FORCED tasks recruit the same parametric MR strategy. To test this, we derived a mental rotation pace parameter (ms/deg) from each task. For the FREE task, this was done by regressing each subjects’ execution trial data (Fig. 4B), and, for the FORCED task, by regressing each subject’s full data set above the critical RT (RT X Movement Angle), as well as fitting a sigmoid to those data (Supplementary Fig. 2). Resulting regression slopes from the FORCED task were then inverted to match the units of the FREE task (ms/deg). Linear MR pace was strikingly similar between tasks (*t*(31) = 0.19, *p* = 0.85; Bayes Factor = 7.81 in favor of the null). Moreover, a significant correlation was found in subjects’ mental rotation paces across tasks (*R^pearson^* = 0.46; *p* = 0.008; *R^spearman^* = 0.41; *p* = 0.02), and this correlation held when using sigmoidal fits on the FORCED data (Supplementary Fig. 2). This result suggests that the same mental rotation computation is operative in both constrained and unconstrained RT contexts.

We note that average RTs in the FREE task (Fig. 4B) were well above those that produced equivalent movement directions in the FORCED task (Fig. 4C). We believe this extra computation time is likely the result of a relatively low sense of urgency in the FREE task compared to the FORCED task: First, we note that the range of RTs observed in the FREE task is comparable to the range observed in a similar study (Georgopoulos & Massey, 1987). Second, the regression analysis on the FREE task (Fig. 4B) revealed no correlation across subjects between the intercept of the regression, which reflects putative non-rotation RT, and its slope, which reflects the mental rotation operation (*R^pearson^* = 0.03; *p* = 0.87; *R^spearman^* = 0.09; *p* = 0.64). This suggests that excess RT in the FREE task is the product of processing unrelated to MR.

#### Control analyses confirm predictions of a parametric-MR strategy

We now address three alternative explanations for the observed linear rise in mean reach directions over RT in the FORCED task (Figs. 4C, D): First, because subjects made many random (i.e. uniformly distributed) movements at low RTs, and gradually made correct movements at higher RTs, a linear trend could appear as an averaging artifact without any truly intermediate movements. However, subjects’ most frequent (mode) reach directions also displayed intermediate values, gradually increasing with RT (Supplementary Fig. 3) and showing a significant linear trend (*p* = 0.002). This argues against this particular averaging confound.

Second, subjects’ non-random reaches could be limited to the 0° target direction (i.e. a “prepotent” response; Stroop, 1935) and the 90° solution direction, with the linear trend merely representing a change in the relative frequency of each as RT increases (another form of averaging artifact). First, we note the mode analysis also argues against this explanation (Supplementary Fig. 3). Second, the full distribution of reach directions plotted in Figure 5A shows no clear evidence for a mode at the 0° location. A subtle degree of bimodality was indeed observed — subjects occasionally reached with an approximately correct magnitude of rotation but with an incorrect sign. We interpret these errors as sign flips, echoing *Experiment 1* (Fig. 3).

**Figure 5.**
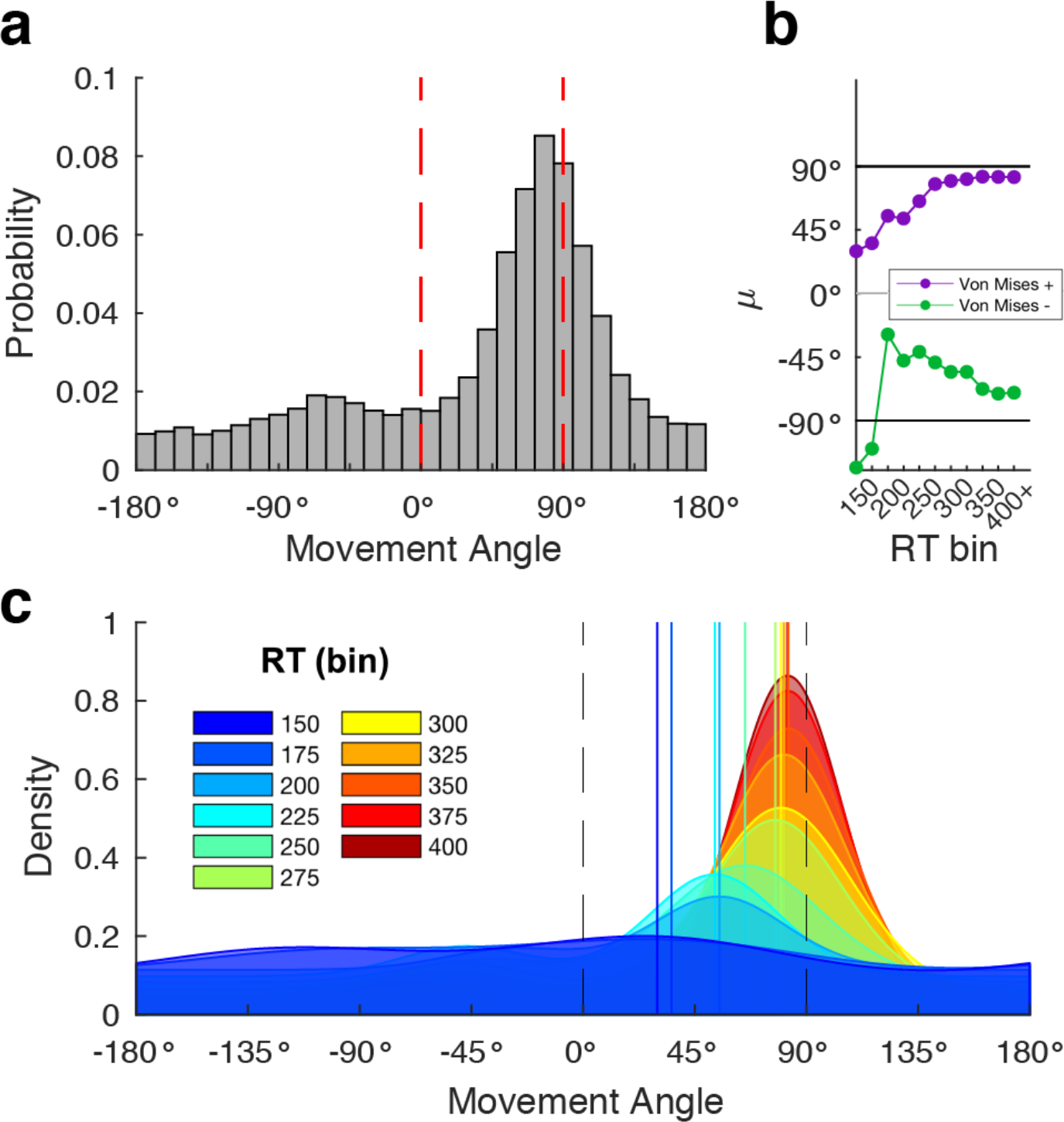
FORCED task full data distribution and model fits. (**A**) Full distribution of movement angles across all RTs in the FORCED task. (**B**) Fitted mean parameters from the “Free-*μ*” mixture model (starting at the 150 ms bin) for directed reaches with a correct sign (Von Mises +, purple) and a flipped sign (Von Mises −, green). See Methods for model details. (**C**) PDFs of model fits in each RT bin. Vertical lines represent the fitted mean parameter (Von Mises +) for each corresponding RT bin.

To further examine the distributions of reach angle over the RT bins, we fit the data with two different mixture models that accounted for random reaches (i.e. modeled with a uniform distribution) and “directed” reaches (i.e. modeled with Von Mises distributions; see Methods for details). The first model, a “Free-*μ*” model, allowed two mean parameters, one positive and one negative, to vary freely over RT bins (thus capturing mental rotation of correct responses and sign flips). The second model, a “Fixed-*μ*” model, fixed the mean parameters at +90° and −90°, which would be consistent with the *a priori* predictions of a discrete RC strategy with swap errors.

As predicted, the mean parameters of the Free-*μ* model showed evidence for MR (Fig. 5B): Both the positive (purple) and negative (green) mean parameters gradually approached, respectively, +90° and −90° as RT increased. Linear regressions on the fit mean parameters revealed a significant positive linear trend for the positive *μ* parameter (*p* < 0.001), consistent with MR. Regression on the negative *μ* parameter, however, revealed no significant trend (*p* = 0.38), though we reasoned this result may be driven by the two deviant points (both > 1.5 sd from the mean) in the first two RT bins (Fig. 5B, green). Figure 5C shows the full probability density functions of the best fit models in each RT bin, with vertical lines representing the positive Von Mises mean in each bin. We emphasize that by accounting for random movements (with a uniform distribution), the mixture model offers further support for the mental rotation of directed movements. Moreover, the apparent rotation of the negative Von Mises mean (Fig. 5B, green) suggests that sign flips themselves undergo MR, albeit it in the erroneous direction. Critically, when the quality of fit of the Free-*μ* model is compared with the Fixed-*μ* model, we find that the Free-*μ* provides a far superior overall fit to the data (ΔAIC = 1189); this large difference comports with MR, as the Free-*μ* model better captures intermediate movements.

Finally, a third alternative interpretation could be that apparent mental rotation would appear as the result of “response substitution” between a prepotent response directed at the target location gradually becoming inhibited, while the rotated response gets excited (Cisek & Scott, 1999). In this interpretation, intermediate movements would be explained by the averaging of simultaneously active motor plans (Cisek & Kalaska, 2005), with one plan directed at the target and one at the solution. Importantly, this model of response substitution and our model of MR make distinct predictions concerning movement speed: Response substitution predicts a slowdown of movement speed on trials where responses are averaged (Aslin and Shea, 1987), whereas MR does not. Movements were fast throughout the FORCED task (*μ* MT = 128.13 ms; Supplementary Fig. 4), and, in fact, movement speed subtly and monotonically increased with RT (Supplementary Fig. 5; *t*-test on regression slopes: *t*(31) = 4.28, *p* < 0.001), arguing against response substitution. The different speed predictions of response substitution versus MR can be captured in the length of a neural population vector, which is correlated with movement speed (Moran & Schwartz, 1999). We conducted a computational modeling analysis using two neural population coding models. These models formalize different movement speed predictions, and provide further evidence that the speed data are compatible with MR but not response substitution (Supplementary Fig. 5). Lastly, we note here that some recent research has questioned the premise that intermediate movements represent involuntary averaging of parallel motor plans (see Discussion; Haith et al., 2015b; Wong and Haith, 2017). However, directly testing these competing hypotheses is beyond the scope of this study.

### Experiment 3: Constrained RT in low set size condition reveals a discrete working memory strategy

*Experiment 3* was designed to provide further evidence for a discrete RC strategy: We used the identical forced-RT task as *Experiment 2*, but with only 2 possible target locations instead of 12, matching the set size manipulation of *Experiment 1* (Fig. 1A). We hypothesized that subjects’ behavior in this context would reveal a bimodal distribution of reaches reflecting cached responses, with no evidence for mental rotation (i.e. intermediate movements). (Note, we did not include a FREE task for the 2-target condition in *Experiment 2*, given that the FREE task itself requires a large variety of rotation magnitudes and signs to test for a linear relationship between rotation and RT; under these conditions, a discrete RC strategy is not plausible given working memory constraints. However, if we constrained it to a single rotation magnitude and sign, it degenerates into the 2T condition of *Experiment* 1.)

With only 2 possible target locations, subjects could fully counteract the rotation with very short RTs (Fig. 6). Examining the circular means of the movement angles (Fig. 6A) reveals an abrupt jump from highly variable movements at short RTs (< 225ms) to consistent movements directed at the rotation solution (90°). The pooled vector plot (Fig. 6B) does not show the several highly-variable intermediate values seen in the circular means (Fig. 6A), suggesting that those are likely the result of circular averaging of extreme values (e.g. −90° and +90°; see below for model fitting). Accordingly, the full histogram of reaching directions (Fig. 7A) is consistent with a bimodal response distribution: Subjects’ responses were concentrated near the solution (+90°) and its opposite (−90°).

**Figure 6:**
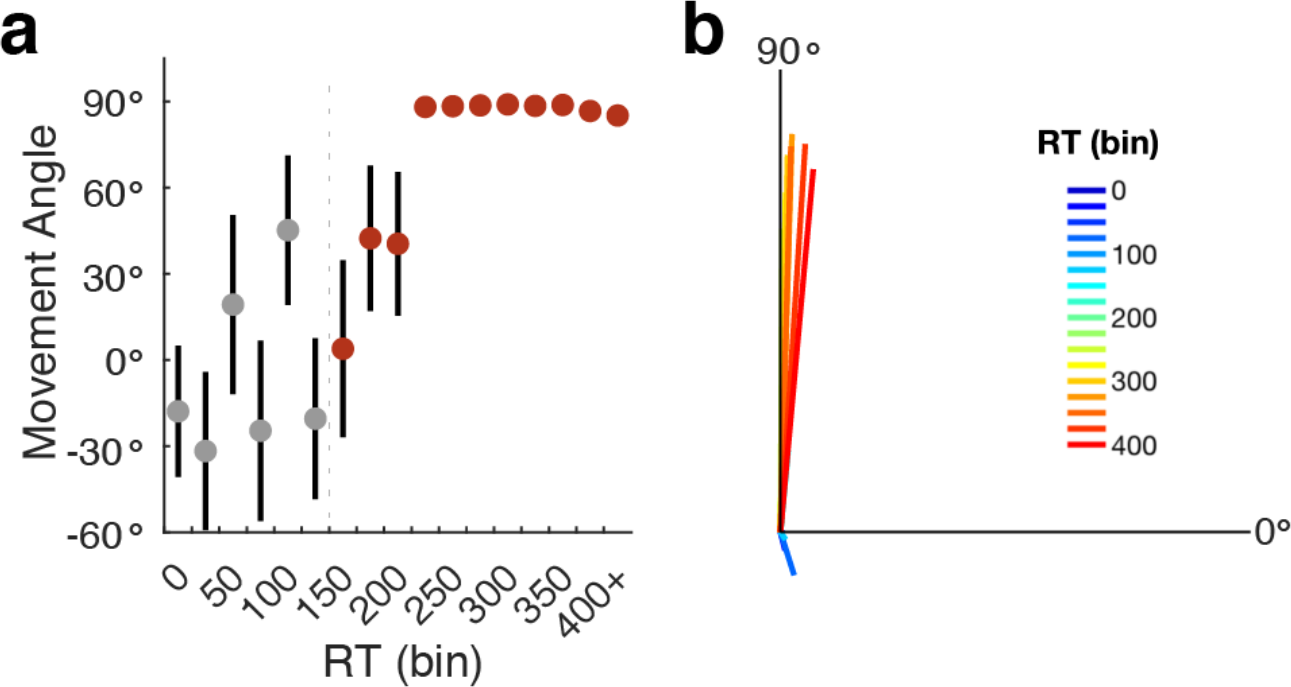
Experiment 3 results. (**A**) Circular mean of movement angles as a function of RT (binned in 25 ms bins) in the 2-target FORCED task. (**B**) Pooled vector representation of data in (**A**). Error bars = 1 s.e.m.

**Figure 7:**
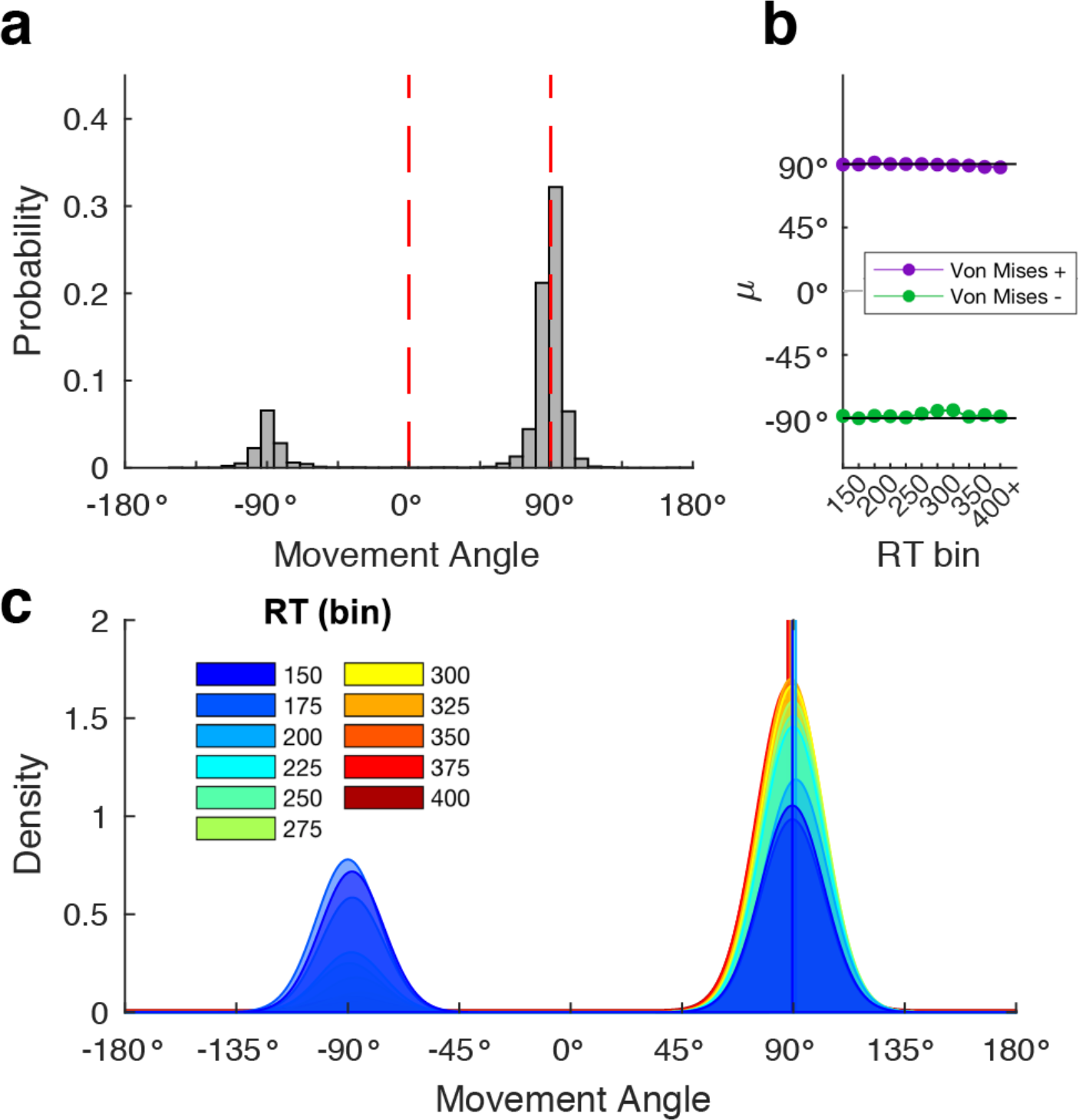
2-target FORCED task full data distribution and model fits. (**A**) Full distribution of movement angles across all RTs in the 2-target variant of the FORCED task. (**B**) Fitted mean parameters from the “Free-*μ*” mixture model (starting at the 150 ms bin) for directed reaches with a correct sign (Von Mises +, purple) and a flipped sign (Von Mises -, green). See Methods for model details. (**C**) PDFs of model fits in each RT bin.

For our next analysis, we again analyzed the mode of pooled movement angles across the RT bins. Unlike the 12-target condition in *Experiment 2*, we did not find a linear trend in the 2- target condition (*p* = 0.20; Supplementary Fig. 3). To better quantify the distribution of reaches in the 2-target condition, we fit the Free-*μ* and Fixed-*μ* mixture models to these data. Critically, the Von Mises mean parameters for the Free-*μ* model appeared to saturate immediately, not displaying the signature of mental rotation seen in *Experiment 2* (Fig. 7B). In fact, linear regression on the positive mean parameter revealed a slight negative slope (*p* = 0.004), which is in the wrong direction for mental rotation, and regression on the negative mean parameter revealed no significant trend (*p* = 0.36). When the quality of fit of the Free-*μ* model is compared with the Fixed-*μ* model, we find that the Fixed-*μ* model provides a far superior fit to the data (ΔAIC = 1941), also in contrast to *Experiment 2*. This large difference comports with a discrete RC strategy, as there is no need for shifting mean free parameters in the absence of intermediate movements, and thus the model is not penalized for unnecessary complexity.

Finally, we sought to directly compare results between the 12-target (*Experiment 2*) and 2-target (*Experiment 3*) forced-RT conditions. First we performed serial *t*-tests between circular mean reaching directions at each RT bin after the critical bin. Means were significantly different at each bin in the 10th - 13th RT bins (all *p’s* < 0.05), spanning the range of RTs between 200 and 300 ms. Next, we performed within-condition *t*-tests at each RT bin to look for significant differences between the group mean movement angles and the solution at 90°. In the 12-target condition, all bins after and including the 7th bin were significantly different from 90° (all *p’s* < 0.01). In the 2-target condition, movement angles were not significantly different from 90° starting at the 8th bin onward (all *p’s* > 0.05). Finally, we performed a more granular model comparison analysis: Recall we found that the 12-target condition (*Experiment 2*) was better fit by the Free-*μ* model, consistent with using a parametric MR strategy, and the 2-target condition (*Experiment 3*) was better fit by the Fixed-*μ* model, consistent with a discrete RC strategy. However, for these analyses we summed AIC values across all model fits after and including the critical (7th) RT bin. Here, we computed ΔAIC between models at each RT bin separately (Fig. 8). The Free-*μ* model fit the 12-target group better at every RT bin (ΔAIC range: 6.92 - 276.13). In contrast, the Fixed- *μ* model fit the 2-target group better at every RT bin (ΔAIC range: 15.69 - 345.46). Thus, consistent with our predictions, subjects in the 12-target condition were likely using a parametric MR strategy whereas subjects in the 2-target condition were likely using a discrete RC strategy.

**Figure 8:**
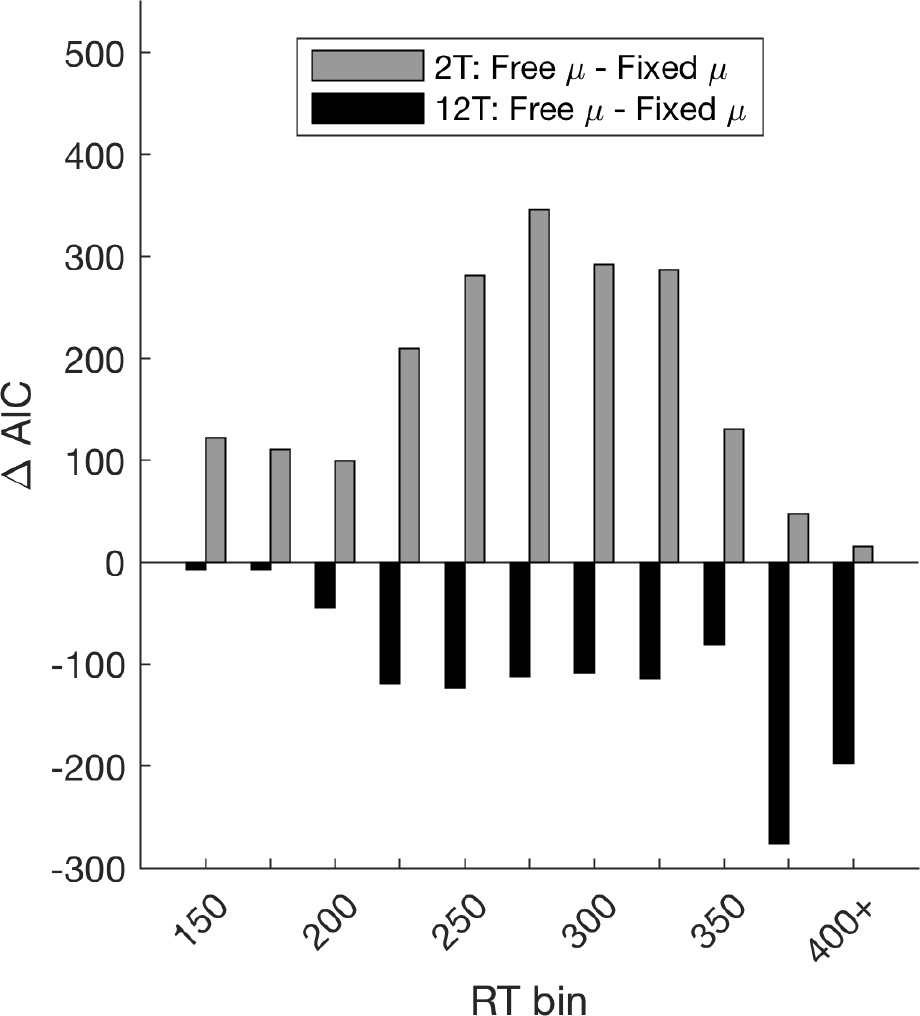
Model comparison. ΔAIC values over each RT bin between the Free-*μ* and Fixed-*μ* models for the 2- target (grey) and 12-target (black) FORCED-RT experiments. Negative values constitute better fits for the Free-*μ* model, and positive values constitute better fits for the Fixed-¼ model.

#### Mental rotation pace from FREE task predicts RT differences during visuomotor learning

By intermixing different rotation sizes, the FREE task in *Experiment 2* provided us with a precise estimate of subjects’ mental rotation paces when RT was unconstrained (Fig. 4B). If our hypothesis is correct, the average mental rotation pace from the FREE task should correspond to observed RT differences between the 75° and 25° conditions from *Experiment 1*. Moreover, this correspondence should hold for the 12T conditions at both the beginning and end of learning, but only the beginning of learning in the 2T conditions in *Experiment 1* (Fig 2C). As shown in Figure 9, the regression line predicted by the FREE task was consistent with MR occurring in all subjects early in learning (Fig. 9A; slope difference between FREE task distribution of slopes and early learning regression slope for 12T groups: *t*(31) = 1.37, *p* = 0.18; 2T groups: *t*(31) = 1.55, *p* = 0.13), but only the 12T groups late in learning (Fig. 9B; FREE task slope difference from 12T groups: *t*(31) = 0.78, *p* = 0.44; 2T groups: *t*(31) = 6.16, *p* < 0.001). This result is consistent with a set-size- dependent shift during learning from an algorithmic working memory strategy to discrete retrieval (Fig. 9C; Provost et al., 2013; Logan, 1988).

**Figure 9:**
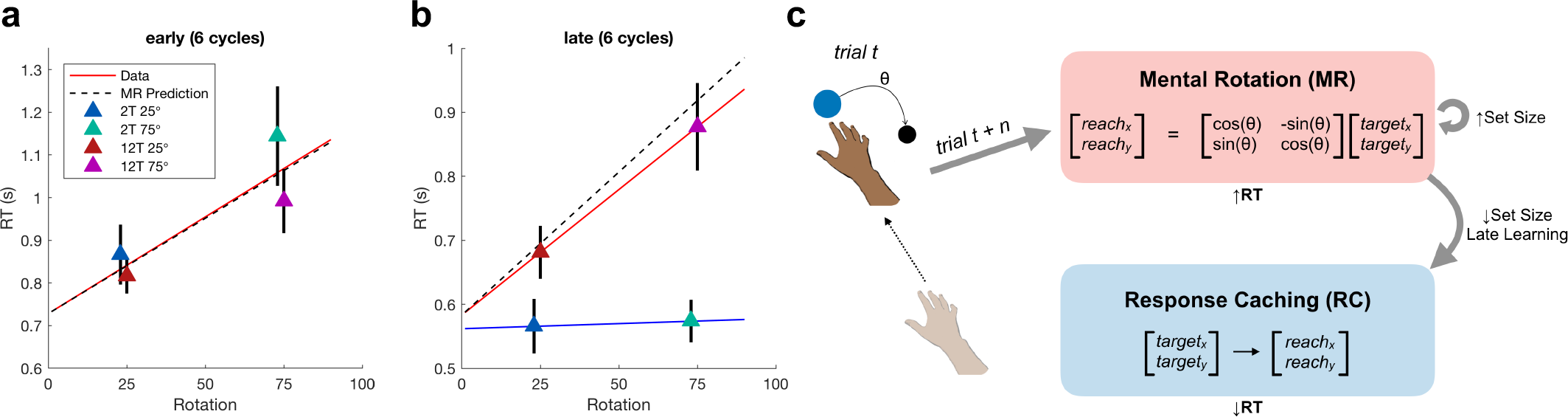
Mental rotation pace from FREE task matches RT data in Experiment 1. (**A**) Mean of median RTs in early learning from *Experiment 1* (triangle markers) with a regression line, for visualization, on all four groups’ data (red), and the regression line predicted by the mental rotation pace gleaned from the FREE task using the same intercept (dashed black). (**B**) Same as (**A**) on late learning data, with separate regression lines for the 12T groups (red) and 2T groups (blue). (**C**) Schematic describing a parametric to discrete learning shift, which is mediated by set size. Error bars = 1 s.e.m.

### Experiment 4: Generalization is affected by learning strategy

How do different working memory representations for visuomotor learning affect generalization when conditions change? In *Experiment 4*, we investigated how different strategies would affect the transfer of visuomotor learning to novel stimuli. We reasoned that subjects using a discrete response caching strategy (RC) would show diminished generalization relative to subjects using a parametric mental rotation strategy (MR). This could occur because under an RC regime, specific local instances are learned, whereas under an MR regime, a global rule (or “structure”) is learned that can be applied indiscriminately.

In *Experiment 4*, subjects were trained on a 45° rotation in a constrained region of the workspace with either 2 targets (2T) or 8 targets (8T), with the width between the furthest targets matched between conditions (Fig 10; see Methods for details). After a brief rotation training block, subjects experienced a generalization test that involved extrapolating their learning to novel target locations (Fig. 10). While this experiment could not infer subjects’ learning strategies in the same way as *Experiments 1-3*, we reasoned that the set size manipulation should bias subjects toward an RC strategy in the 2T condition and an MR strategy in the 8T condition.

**Figure 10:**
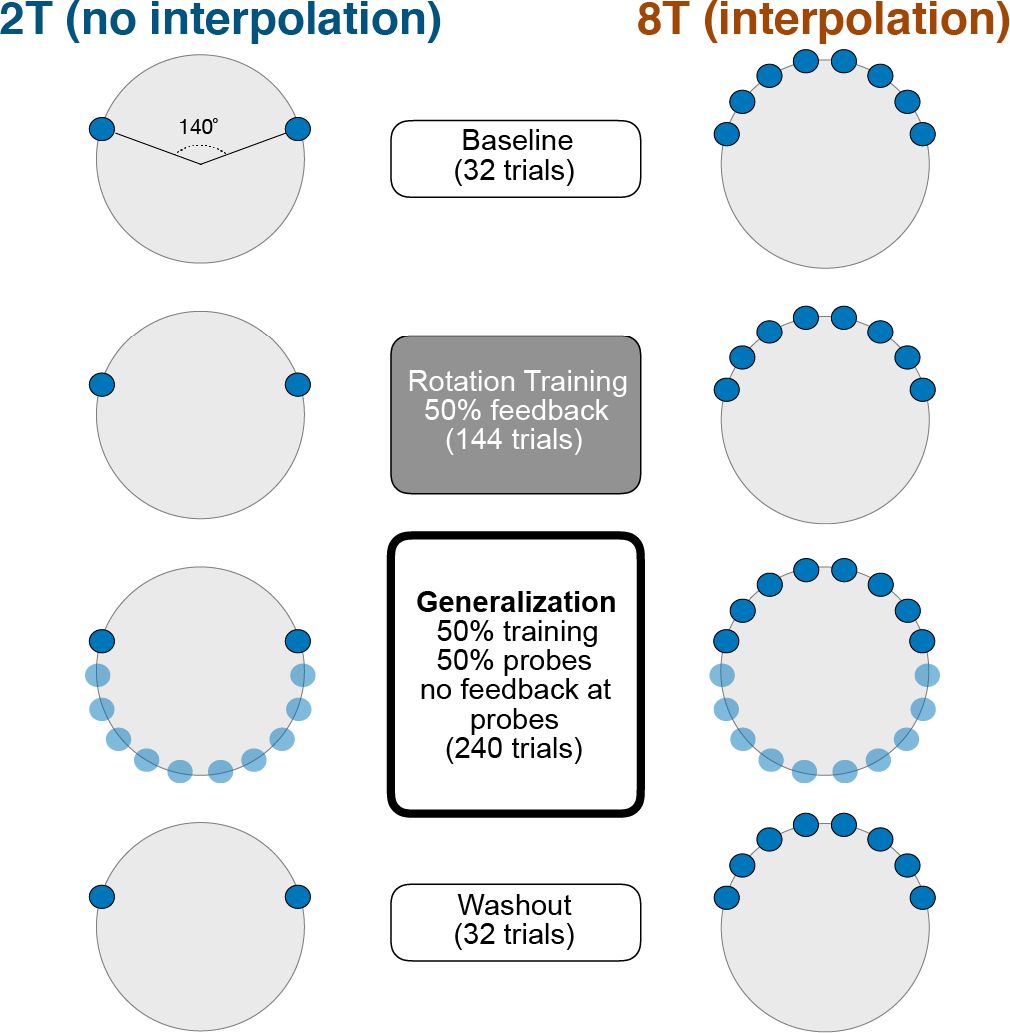
Experiment 4 design. Subjects learned to counter a 45° rotation in either a 2T learning condition (left) or an 8T learning condition (right). In a generalization block, novel targets were presented.

As predicted, subjects in the 2T group showed more narrow generalization relative to the 8T group (Fig. 11A). These effects were generally symmetric, as shown in Figure 11B. To investigate group differences, we performed a trial-by-trial regression analysis on subjects’ movement angles toward the generalization targets (Fig. 11C; Fig. S6; see Methods for details). First, we found that the amount of practice (i.e., a trial-number regressor) predicted an increase in movement angles to the generalization targets for both the 2T (*t*(14) = 2.73, *p* = 0.02) and 8T groups (*t*(16) = 2.35, *p* = 0.03), suggesting that as the generalization block progressed, generalization increased.

**Figure 11.**
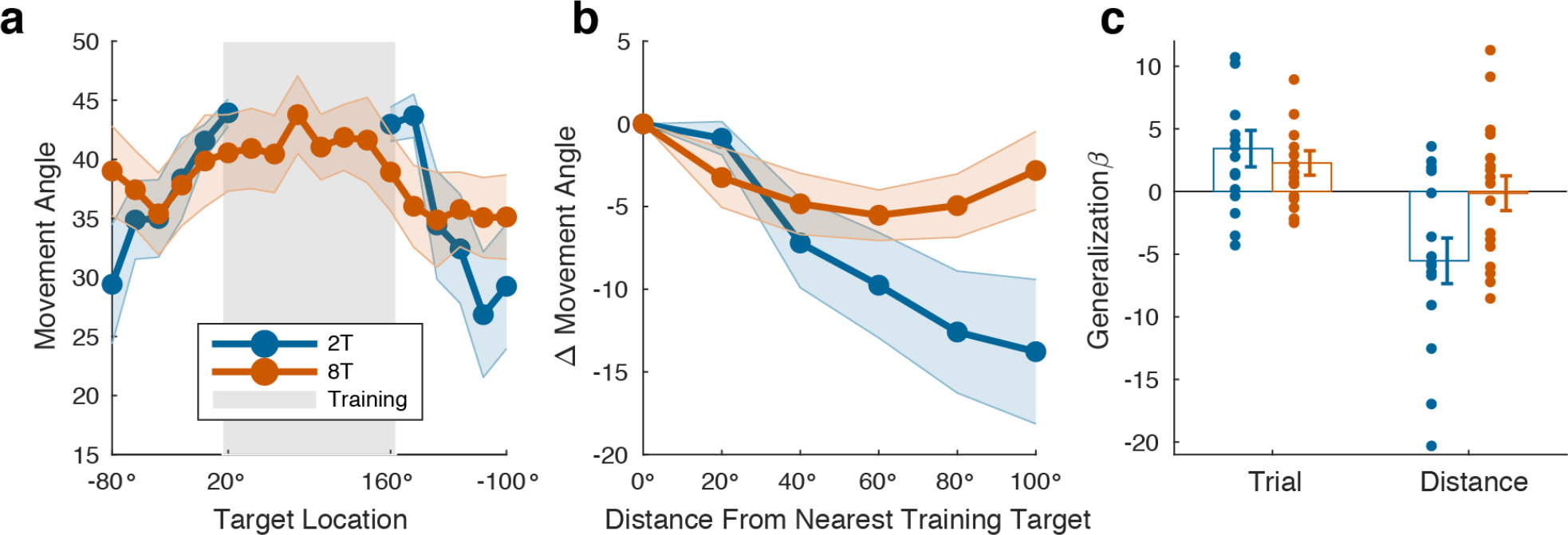
Experiment 4 results. (**A**) Mean movement angles in the generalization block targets and training block targets (gray shading). (**B**) Mean change in movement angle relative to mean movement angles at the nearest training targets. (**C**) Results of the regression analysis, showing effects of trial number and generalization target distance on movement angles in the generalization block. Shading and error bars represent 1 s.e.m.

Consistent with our main hypothesis, we found that for the 2T group the distance of generalization targets from the nearest training target negatively impacted movement angles (*t*(14) = 3.06, *p* = 0.009). This same effect, however, was not significant in the 8T group (*t*(16) = 0.10, *p* = 0.92). Critically, a two-sample *t*-test on regression weights between the two groups revealed a significant difference (*t*(30) = 2.41, *p* = 0.02). Thus, the working memory representation recruited for explicit visuomotor learning appears to shape how the newly learned behavioral policy is generalized.

## DISCUSSION

The role of working memory in motor learning is not well understood, though it is clear that controlled, deliberative processes are important for learning (Taylor et al., 2014; McDougle et al., 2016; Fernandez-Ruiz et al., 2011; Huberdeau et al., 2017). Here we provide evidence for two working memory strategies that contribute to learning in a visuomotor rotation task — discrete response caching (RC) and parametric mental rotation (MR). A parametric MR strategy manifests as a linear relationship between rotation magnitude and RT, consistent with classic mental rotation (Shepard and Metzler, 1971; Georgopoulos and Massey, 1987; Bhat and Sanes, 1998). In contrast, a discrete RC strategy manifests as a look-up table of S-R relationships (Petrides, 1985; Collins and Frank, 2012), consistent with capacity-limited working memory (Cowan, 2010; Luck & Vogel, 2013).

How do parametric versus discrete working memory representations relate to long term skill acquisition? One useful analogy here could be the dissociation of model-based and model-free reinforcement learning (Daw et al., 2005), where the former relies on an explicit model of transition probabilities between responses and sensory states, and the latter merely reinforces rewarded actions. One speculation could be that model-based computations, which could be analogous to parametric motor learning strategies, are themselves made automatic over time (Economides et al., 2015). This would suggest that the transition from parametric to discrete strategies we observed (Fig. 9) could represent mental rotation itself becoming automatic. However, if a complex computation like mental rotation becomes “cached,” it is hard to discern if the computation can still be said to be operative — it could be that the same responses that reflect the computation have been cached, and the computation itself is thus no longer needed (Logan, 1988; Haith and Krakauer, 2018). This kind of caching could represent an intermediate form of processing that lies somewhere between model-based planning and model-free habit.

One idea could be that during learning, there is a first transition from model-based planning (e.g. leveraging a model of the world) to retrieval-based S-R (Fig. 9C). This form of S-R could then itself be proceduralized as model-free habit, no longer requiring working memory. In this framework, difficult computations are not transformed or discarded during skill learning, but rather come to be bypassed during response preparation. Future studies using extended training could test this hypothesis. However, we note that our observation of swap errors (Fig. 3) suggests that S-R learning in our task had not been fully automatized and still likely involved working memory (Bays, 2016). Extending training over days or weeks can better assess the proceduralization of learning (Huberdeau et al., 2017).

Presumably, it is important that an algorithmic strategy can still be flexibly recruited as needed after training, especially if the environment changes. Several studies on long-term visual mental rotation (using the task originally described in Shepard and Metzler, 1971) find that although practicing mental rotation for weeks leads to an exponential reduction in overall RT, the mental rotation effect (i.e. RT as a linear function of rotation magnitude) diminishes only subtly and may not go away (Leone et al., 1993; Wright et al., 2008). In the longest version of this task (Leone et al., 1993), a set size of 12 different objects and many rotation magnitudes was used, thus making a discrete retrieval strategy onerous. In a different study with long-term practice of visual mental rotation, only three stimuli and four rotation angles were used, and mental rotation effects significantly decreased after extensive training (Tarr and Pinker, 1989). Intriguingly, when novel orientations were introduced at the end of learning, the mental rotation effects returned, suggesting a “dual process” model where subjects either retrieve an over-learned object from rote memory or perform a parametric computation (Tarr and Pinker, 1989). This model was confirmed in a recent study by Provost et al. (2013), which showed that repeatedly performing mental rotation on a small number of objects leads to the disappearance of mental rotation RT effects after days of training; however, when a large number of items are used during training, the RT effects persist. These results in the visual domain overlap considerably with our results in the motor domain, suggesting homologous mechanisms of spatial cognition. Moreover, these effects offer strong empirical support for Logan’s (1988) theory of automaticity, where he hypothesized that skill learning involves a transition from algorithmic to retrieval-based strategies.

We note that the results of our 12-target forced-reaction-time task (Fig. 4C, D) also have broad implications for the study of mental imagery. Our results provide novel evidence supporting a critical prediction of analog cognitive computations, namely that mental rotation proceeds through intermediate states (Shepard and Metzler, 1971). These intermediate states have been implied from RT measures (Cooper, 1976) or neural population statistics (Georgopoulos et al., 1989), but have not been rendered in overt motor behavior. Our results suggest that a mental rotation-like operation can drive volitional re-planning of a movement goal via a continuous sweep through direction space at a constrained pace. We note that this mechanism doesn’t require that neurons necessarily be directionally-tuned, but could instead represent a low-dimensional projection of a high-dimensional representation onto the 2D angular space prescribed by our task (Churchland et al. 2012). The aforementioned results in motor cortex (Georgopoulos et al., 1989) were challenged by an alternative explanation, which posits that the gradual “averaging” of an initial motor plan at 0° with a second plan at 90° could give the appearance of mental rotation (Cisek & Scott, 1999). This response substitution account has been supported by the behavioral finding that subjects appear to involuntarily average co-active motor plans in “go-before-you-know” paradigms (Stewart et al., 2014) or “target-jump” saccade tasks (Aslin and Shea, 1987), where goals are cued late in a trial.

This averaging hypothesis has been challenged by recent findings suggesting that nominally averaged plans may actually be the product of a decision-making process optimizing a single movement plan to account for multiple goals (Haith et al., 2015b; Wong and Haith, 2017). In a recent study, Wong & Haith (2017) used a go-before-you-know task where subjects had to initiate a movement between two targets before one was cued as the goal. Critically, when moving relatively slowly (average movement times ~ 600 ms), subjects often reached at an angle between two competing targets until the goal target was cued, after which they swerved to the correct choice. However, when moving quickly (average movement times ~ 300 ms), subjects often moved in a straight line to one of the two options, making no corrections after the cue. These and other similar results (Haith et al., 2015b) suggest that when making reaching movements between two competing options, subjects’ movement directions may reflect an optimal decision process that can buy itself time to make the correct decision under uncertainty. Critically, not only did our FORCED task have a single goal (a 90° reach) instead of multiple goals, but, more importantly, subjects elicited rapid reaching movements (average movement times ~ 130 ms) well below the movement time threshold for plan averaging (Wong & Haith, 2017). Taken together, our task design and results suggest that neither obligatory averaging of co-active motor plans (Stewart et al., 2014) nor strategic plan averaging (Wong & Haith, 2017) describe our results. We maintain that parametric rotation of a motor plan is the most parsimonious explanation.

One trend in our data is the tendency for subjects to “under-rotate” when they were putatively using a parametric MR strategy. This can be observed in *Experiment 1* (12T groups; Fig. 2B), both the FREE and FORCED tasks of *Experiment 2* (Fig. 4), and the 8T group’s training target behavior in *Experiment 4* (Fig. 11A). In contrast, when performing discrete RC, subjects appear to reach all the way to the solution (2T groups Fig. 2B; Fig. 6). We do not have a clear result that speaks to this issue. One speculation is that due to the extra computation time needed for mental rotation, a latent “urgency” signal may drive subjects to initiate their movement before they’re finished rotating, perhaps reflecting a speed-accuracy trade off. Another detail in our data is that MR in the 12-target FORCED task (*Experiment 2*) appears to start with a rapid jump followed by a linear rotation; this is most clearly shown when looking at the model fit (Fig.5B). This suggests that the MR process may involve a complex sequence of discrete rotations with varying magnitudes.

In terms of the neural substrates of parametric MR and discrete RC, we present several hypotheses. For parametric MR, parietal areas — the putative locus of mental rotation and similar sensorimotor transformations (Kosslyn et al.,1998; Harris & Miniussim, 2003; Bueno & Andersen, 2006; Zacks, 2008) — could continually feed a shifting movement goal to motor cortex, which would explain the observed intermediate movements when RT was cut short. Consistent with this hypothesis, Anguera and colleagues (2010) found overlapping activation in inferior parietal and dorso-lateral prefrontal cortices during both a visuomotor adaptation task and a mental rotation task. Discrete RC, on the other hand, being a form of capacity-limited item-based working memory, could rely on the maintenance of S-R relationships in dorso-lateral prefrontal cortex (Curtis & D’Esposito, 2003). Future studies could test these hypotheses using brain imaging or stimulation techniques.

A dissociation between parametric and discrete memory representations is a theme in several learning and memory domains. One example is human mathematical reasoning, where there are strong behavioral and neural dissociations between numerical computations learned by rote and then recalled from semantic memory (“two times four is eight”), versus flexible parametric operations performed on a “mental number line,” represented in fronto-pareital spatial working memory regions (Deheane and Cohen, 1997; Hubbard et al., 2005). Here we present an analogous dissociation, showing evidence for both the maintenance of discrete motor responses, and the parametric manipulation of such responses, in working memory. Movement plans may be held and manipulated in working memory, perhaps in a manner consistent with visual, auditory, and tactile representations. Overall, the results presented here suggest that spatial cognition occupies a key role in motor learning.

## METHOD

### Participants

158 right-handed subjects (age range 18-34, 73 women) were recruited from the research participation pool maintained by Princeton University in exchange for course credit or monetary compensation. Handedness was verified using the Edinburgh handedness inventory (Oldfield, 1971). All subjects participated in accordance with the university’s IRB and provided written, informed consent. In *Experiment 1* (n = 80), 20 subjects were used per condition. A single subject was excluded for disregarding the key task instruction of attempting to land the cursor on the target (asymptotic movement error > 6 sd from the mean). We note that the sample size was not determined by a power analysis, although it is consistent with sample sizes in other studies using similar tasks (Stewart et al., 2014; Wong and Haith, 2017; McDougle et al., 2017). In *Experiment 2*, a power analysis was used (alpha = 0.95) and revealed that a sample of 19 subjects would replicate the effect size of a relevant correlation result (*d* = 0.66; correlation of mental rotation paces in Pellizzer and Georgopoulos, 1993). To be conservative, we sought to approximately double it, so we recruited a sample size of 32 (given counterbalancing). This is also in the range of a previous study that used a similar within-subject design and a somewhat similar task, which had a sample size of 26 (Pellizzer & Georgopoulos, 1993). A power analysis was used to determine the sample size in *Experiment 3*, which aimed to test the regression effect seen in *Experiment 2 (d* = 2.60; regression on movement angle X RT). The necessary sample size was 5, but, again to be conservative and counterbalance rotation signs, we recruited a sample of 10. For *Experiment 4*, we did not have a salient comparable analysis with which to conduct an *a priori* power analysis (i.e. regression weights on generalization probe distance versus movement); however, in a previous study where we modeled generalization (McDougle et al., 2017), we used a sample of 15 subjects. Thus, we opted to use 36 subjects in total (18 per group), a number which allowed for symmetric counterbalancing according to the task design.

### General Experimental Procedures and Analysis

In all experiments, subjects made rapid, center-out, open-loop reaching movements to visual targets (5.0 mm radius) using a digitizing tablet, holding on to a digital pen with their hand in a power grip position and sliding the pen across the tablet (Intuous Pro; Wacom). The task was controlled by custom software written in MATLAB (Mathworks, Natick, MA; Psychophysics Toolbox). Hand position was sampled at 140 Hz. Stimuli were shown on a 21.5-inch LCD computer monitor (Planar), mounted horizontally 25 cm above the tablet, occluding vision of the hand. A small cursor (2.5 mm radius) provided endpoint feedback after each reach terminated. Analyses were conducted in MATLAB and R (GNU).

### Experiment 1

Subjects (N = 80) made rapid, 7 cm movements to targets in a blocked reaching task using a 2 X 2 between-subjects design, crossing the factors rotation magnitude and number of targets (“set size”; Fig. 1A). For the rotation, two magnitudes were used, ±25° and ±75°, with the sign of the rotation counterbalanced within each condition. For set size, subjects were exposed to either two targets (2T) or 12 targets (12T). In the 12T condition, target locations were pseudorandomly presented at 12 possible locations (0°, 30°, 60°, 90°, 120°, 150°, 180°, 210°, 240°, 270°, 300°, 330°), where the same location was never repeated in consecutive trials. In the 2T condition, two targets were presented, and the locations were randomized across subjects to include one position from the 12 possible locations above and its opposite 180° away (Fig. 1A). Within a subject, targets were presented pseudorandomly, where the same target was not repeated more than twice in a row. At the start of every trial, to position their hand in a central “start” position (5 mm radius) subjects used a dynamic visual ring that was scaled based on their distance from the start. After holding the start for 500 ms, a visual target appeared and cursor feedback was removed.

In a baseline block (36 trials), subjects reached to the targets with veridical endpoint feedback. At the start of the subsequent rotation block, the rotation was abruptly applied and was maintained for 300 trials, all with endpoint feedback. Finally, in the washout block, subjects were told to cease any strategy they may have adopted to counter the rotation and reach directly for each target. At the start of the experiment, subjects were told to do their best to land the cursor on the target. If the cursor hit the target, a pleasant chime was sounded, otherwise an aversive buzzer was sounded. In the washout block a neutral sound was played after each reach. Critically, to limit implicit sensorimotor adaptation and better isolate strategic explicit learning, all feedback was delayed by 2 s on every trial (Kitazawa et al., 1995; Brudner et al., 2016; Schween et al., 2017; Parvin et al., 2018).

Movement angles were computed as the angle of the hand, relative to the target, when it crossed the invisible target ring of 7cm. All movement angles were rotated to a common reference frame about the 0° axis of the unit circle. In all experiments, reaction time (RT) was measured as the time elapsed from target appearance to the hand leaving the start circle (i.e. 5mm from the center of the tablet), and movement time (MT) was measured as the duration of the movement from the end of the RT to the time at which the hand crossed the invisible workspace ring. Lastly, trials where RTs exceeded three standard deviations above or below a subject’s mean RT (1.30% of trials), or trials where movement angles exceeded three standard deviations above or below a subject’s mean movement angle (0.87% of trials) were excluded from analyses (the latter exclusions were not performed for the sign error analysis in Fig. 3).

### Experiment 2

Subjects (N = 32) performed two tasks (FREE and FORCED; Fig. 1B), with the order of completion counterbalanced, as well as the rotation sign used in the FORCED task (−90° versus 90°). The FREE task was designed to verify the classic signature of mental rotation, and the FORCED task was designed to interrupt mental rotation.

### FREE Task

At the start of every trial, subjects used continuous cursor feedback to position their hand in a central “start” position (5 mm radius). After holding for 400 ms, a visual target appeared and cursor feedback was removed. Participants were instructed to make a rapid, straight shooting movement to the target. The trial concluded when the hand crossed an invisible ring (8 cm radius), at which point feedback of the cursor was provided at the location where the hand crossed the ring. Slow movement times (MT) were discouraged: If MT exceeded a 700ms limit, a “too slow” warning was delivered visually to the subject by the task software. In line with the instructions, subjects moved straight and rapidly, with a mean MT of 208.32 ± 8.54 ms (standard error of the mean).

Subjects were instructed that they would be performing a number of trial “pairs” (Fig. 1B, top): In the first trial of each pair, the “learning” trial, subjects were instructed to reach directly at the displayed target and observe where the feedback cursor landed. In learning trials, the target was blue and appeared in one of four off-cardinal locations (10°, 100°, 190°, 280°). In the second trial of the pair, the “execution” trial, subjects were told to apply what they learned about the relationship between their movement and the resultant feedback and attempt to make the cursor terminate within the target. In execution trials, the target was red and appeared in one of the three locations in which the learning target did not appear. Target locations were pseudo-randomized within and between trial pairs. Subjects performed 140 trial pairs. This task was modeled after a previous study (Pellizzer & Georgopoulos, 1993), but with the distinction that subjects were not provided with an explicit symbolic cue regarding the exact solution to the rotation; instead, subjects had to determine the rotation’s size and sign themselves, so that our task echoed canonical visuomotor adaptation paradigms (Krakauer, 2009).

Rotations used in the learning trials ranged from −90° to 90° by 15° intervals. Rotations were pseudo-randomized throughout the task, and each rotation was seen on 10 different trial pairs, except for the 0° null rotation, which was seen on 20 trial pairs. Each subject received an individualized schedule of rotation magnitudes and target locations.

Analyses were conducted on execution trials only, limited to trials where there was an imposed rotation. Movement angles were computed as the angle of the hand, relative to the target, when it crossed the invisible target ring. All movement angles were rotated to the same 0° target axis and matched to a single sign for analysis (see *experiment* 1). The first analysis was a regression on subjects’ reaching angles, relative to the imposed rotation, on execution trials. We chose to use the absolute reaching angle since on a significant number of trials (8.71%), subjects approximated the magnitude of the rotation but misinterpreted the sign (i.e. were within 15° of the “flipped” solution). No significant difference in mean RTs was found between correct and “flipped” execution trials (*t*(31) = −0.99, *p* = 0.33), suggesting that subjects did not hesitate and change their mind on flipped trials, but simply misremembered the proper clockwise/counterclockwise direction at the onset of the execution trial (see Results and Discussion for a further discussion of sign errors).

A one-sample *t*-test was performed on the fitted slopes resulting from the regression of rotation magnitude and movement angle to test for the presence of a reliable trend. The second analysis was a regression of each subject’s realized movement angles onto corresponding RTs. A one-sample *t*-test was performed on the fitted slope values to test for the presence of a reliable trend. Moreover, each subject’s slope derived from this regression analysis served as their mental rotation “pace”, in the units of milliseconds per degree (Pellizzer & Georgopoulos, 1993). We also performed a similar linear regression using the imposed rotation angles as the predictor variable to confirm that it echoed the regression using the actually-realized movement angles; both analyses yielded comparable results, though the movement angle regression is likely to be a better estimate of a subject’s true mental rotation pace (Georgopoulos and Massey, 1987).

### FORCED Task

The FORCED task utilized a modified forced-response-time paradigm to interrupt putative mental rotation (Ghez et al., 1997; Haith et al., 2016; Fig. 1B, bottom). Like the FREE task, at the start of every trial subjects used continuous cursor feedback to position their hand in a central “start” position. Once subjects were positioned in the start, a countdown of four tones was emitted from the computer speakers (Logitech). The tones were played 600 ms apart, and subjects were instructed to try and synchronize the initiation of their reach with the fourth tone, effectively “replacing” that tone. The experiment and instructions were deliberately designed to emphasize early movements over late ones: If the subject initiated their movement > 100 ms after the fourth tone, the screen was blanked and a “Too Late” message was displayed. Subjects could initiate their movement any time after target appearance, to encourage early RTs. However, if they moved before the target appeared the screen was blanked and a “Too Early” message was displayed. Similar versions of this forced-response-time task have been previously used to study the effect of restricted RTs on visuomotor learning and movement preparation (Ghez et al., 1997; Haith et al., 2016).

Subjects were instructed to prioritize reaching on time, and, secondarily, to try and land the cursor on the displayed target. On each trial, the targets could appear in one of 12 off-cardinal locations (10°, 40°, 70°, 100°, 130°, 160°, 190°, 220°, 250°, 280°, 310°, 340°). Endpoint cursor feedback was presented after the subjects passed the invisible target ring, and the appearance of the feedback was delayed by 500 ms. Like *experiment* 1, this brief delay was added to inhibit implicit adaptation to the imposed rotation and thus isolate the cognitive re-aiming process (Kitazawa et al., 1995; Brudner et al., 2016; Schween et al., 2017). A briefer delay was used in this experiment because more trials were desired, and 500 ms delays have been shown to significantly attenuate implicit adaptation (Kitazawa et al., 1995). The delay manipulation was successful, yielding subtle, though significant aftereffects (*μ* = 1.98°; *t*(31) = 2.06, *p* = 0.048).

The moment of target appearance was titrated such that subjects had varying amounts of time with which to compute the target location, plan, and execute their movements. These windows reflected the time elapsed between target appearance and the fixed moment of the fourth tone. Seventeen test RT windows were used, ranging from 200 - 600 ms by 25 ms intervals. An asymptotic RT window was also used, giving subjects up to 1200 ms to react. We reiterate that because subjects could move early in our variant of this task (i.e. any time after target appearance), the windows acted as “guides” rather than being perfectly predictive of subjects’ realized RTs (i.e. subjects tended to move before they needed to; see Results). Finally, catch trials of 0 ms were used to ensure that subjects stayed on task and executed movements on time even if they had not perceived the target yet.

Subjects performed the FORCED task in three blocks. To get accustomed to the task, in the first block subjects performed 64 baseline trials with veridical endpoint cursor feedback, with pseudo-randomized RT windows and target locations. In the subsequent rotation block, subjects performed 624 trials, with 35 trials at each of the test RT windows, 14 trials at the asymptotic RT window, and 15 catch trials. For the rotation block, a fixed rotation of 90° (or −90°, for counterbalancing) was imposed on the cursor. Subjects were thoroughly educated about the rotation before this block began. Subjects were told to try and counter the rotation and land the cursor on the target every trial. Due to the difficulty of the task, subjects were encouraged by a monetary bonus of up to $5 based on their performance in the rotation block, and were informed that missed trials (reaching too late or too slowly) would count against their performance score. Finally, in the aftereffect block, subjects performed 48 trials with a 1200 ms RT window, pseudorandomized target locations, and no cursor feedback. Subjects were told to reach directly for the target on every trial of the aftereffect block. These data were later compared to the baseline data to get an estimate of any implicit adaptation.

All analyses were conducted on trials where subjects reached on time (78.90% of trials). Like *Experiment 1* and the FREE task, movement angles were computed as the angle of the hand relative to the target when it crossed the invisible target ring. Importantly, subjects followed the instructions and made straight shooting movements: Movement times were rapid (*μ* = 128.13 ms) and no feedback was provided during the reach, which helped to ensure movements were straight. Movement speed was computed by taking the average of the derivative of hand position from the time subjects left the start circle to the time they crossed the invisible target ring. (Due to the rapid “shooting” movement required by the task, average speed was used instead of peak speed to reduce noise in the estimate as subjects often reached peak speed after passing the target; we note that this particular approximation of speed did not influence the main results of the analyses related to movement speed).

Our first analysis involved investigating subjects’ reach angles as a function of RT. RT bins were taken every 25 ms, from 0 ms through 400 ms, with the final bin including all RTs above 400 ms. Similar to previous results (Haith et al., 2016), in catch trials (0 ms RT window) subjects prioritized the timing demands of the task when no target appeared and moved relatively randomly around the circle, with some slight biases. A single “critical RT bin” was computed, after which reaches were determined to be primarily non-random (Supplementary Fig. 1). Linear regressions were performed on subjects’ full distribution (i.e. no binning) of reach angles and RTs after the critical RT bin, and a *t*-test was performed on the resulting slopes to test for the presence of a significant trend. These slopes were used as the mental rotation “pace” parameters in further analyses.

Mental rotation paces were compared between the FREE and FORCED tasks in four ways: First, a one-sample *t*-test was performed to show that the null could not be rejected.

Second, a Bayes Factor was computed on the resulting one-sample t-value using the JZS method (Rouder et al., 2009) to quantify evidence for the null. Third, both parametric (Pearson) and non-parametric (Spearman) correlations were conducted between the values to test for a significant relationship that is robust to outliers (both metrics were used for all correlations in this study). To confirm the robustness of this correlation, a secondary analysis that involved fitting a sigmoid to the data was used as an alternative method for extracting the pace parameter in the FORCED task. Lastly, two supplementary modeling analyses were conducted to test alternative interpretations, involving both a mixture model analysis (see below; Fig. 5), and a neural model which modeled movement speed and direction using a cosine-tuned population coding model (Supplementary Fig. 5).

### Experiment 3

Subjects (N = 10) performed the FORCED task used in *Experiment 2* (Fig. 1B), with one critical difference: Only two target locations were used, where one was drawn from one of the 12 possible locations of *Experiment 2* (FORCED), and the other target was its 180° counterpart. The particular pair of targets used, and the sign of the 90° rotation, were counterbalanced across subjects. All task instructions and analyses matched those described in *Experiment 2*. MTs were similarly rapid (*µ* = 111.25 ms), and aftereffects were similarly small (*μ* = 3.17°) but did not reach significance (*t*(9) = 2.12, *p* = 0.06). For comparison purposes, the same “critical” RT bin was used in this experiment as that derived from *Experiment 2* (RTs > 150 ms; see above).

### Experiment 4

Subjects (N = 36) performed a reaching task (Fig. 10) that was identical to *Experiment 1* in terms of basic trial design, visual stimuli, and feedback timing. In a baseline block (32 trials), subjects reached to visual targets with veridical feedback. In the rotation block (144 trials), subjects experienced a consistent 45° rotation (or −45°, for counterbalancing), and received cursor feedback on 50% of those trials. In the learning block, subjects were divided into two groups: a 2T group and an 8T group. In the 2T group, learning targets appeared at one of two locations 140° apart, with the specific pair of locations counterbalanced across subjects. In the 8T group, learning targets appeared at one of 8 possible locations, spaced equally in a 140° region, with the specific locations also counterbalanced across subjects.

The generalization block tested how subjects extrapolated their learning to a new region of space. On 50% of trials, targets appeared at one of the previously-presented target locations, and in the other 50% of trials targets appeared at one of 10 equally-spaced novel locations within the 220° region of the workspace lying outside of the 140° training region. To test generalization without allowing for new learning at the generalization targets, no feedback was given on generalization probe trials. However, feedback was shown on all trials where a previously-seen learning target was presented. Thus, learning at the original training locations was maintained, and the overall probability of seeing feedback was matched between the generalization and learning blocks (the latter was done to limit the “context change” brought about by the generalization block). In the generalization block, subjects were instructed to reach to novel targets in a manner that would cause the cursor to hit the target if they had seen the feedback. In a final washout block, subjects were told to cease any strategy they were using to counter the rotation and reach directly for the presented targets.

Movement angles were computed in the same manner as *Experiment 1*. Given the limited number of feedback trials in the rotation training block (72 trials), we first analyzed whether subjects successfully adopted a strategy to counter the rotation in this brief period of time and with the added interference of no-feedback trials. We used an *a priori* learning criteria derived from a recent paper (Poh et al., 2016): In each subject, a one-sample *t*-test was performed against 0° on movement angles over the last 4 trial cycles (optimal movement angle = 45°). Four out of thirty six subjects showed non-significant asymptotic learning (*p* > 0.05), and were thus excluded from the generalization analysis. Following *Experiment 1*, trials where RTs exceeded three standard deviations above or below a subject’s mean RT, or movement angles exceeded three standard deviations above or below a subject’s mean movement angle, were excluded (1.15% of trials).

Generalization was analyzed as follows: First, subjects’ movement angles were rotated to a common reference frame so that the learning target region lied between 20° and 160°. For visualization purposes, movement angle generalization functions were computed according to both the raw target angle (Fig. 11A) and the change in movement angle as a function of the target’s absolute distance from the nearest learning target (Fig. 11B). For group comparisons, a trial-by-trial regression analysis was performed using movement angles on generalization trials (i.e. movements to novel targets) as the dependent variable, and four separate z-scored regressors: The trial number, the distance of the current target from the nearest learning target, the subject’s RT, and the interaction of RT and distance. (Non-distance related regressors were added to better isolate the main effect of target distance.) Two-sample *t*-tests were performed on the resulting beta values of interest (Fig. 11C).

For completeness, we also conducted a traditional generalization analysis, where Gaussian functions are fit to each subject’s generalization data (McDougle et al., 2017). However, because of nearly complete generalization in many subjects (especially in the 8T group), various combinations of the height, width, and offset free parameters can yield flat generalization functions, leading to unstable parameter estimation (see Fig. S6 for details). Thus, we opted for the more interpretable regression approach.

### Mixture Model

A mixture model was used to characterize data in the FORCED tasks from *Experiments* and *3*. In this analysis, we modeled reach data as a mixture of two circular normal (Von Mises) distributions representing both positive (correct) and negative (flipped) directed reaches, and a single uniform distribution that represented random reaches. The model had probability density function,

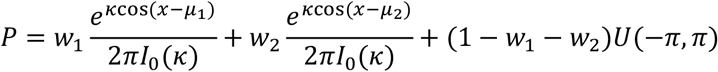

where *I_o_* is the modified Bessel function of order 0, *w* is the weight given to each distribution (*w*_1_ + *W*_2_ ≤ 1), *κ* is the concentration parameter (which we fit as a single parameter between the two Von Mises pdfs), and *U*(−π, π) is the uniform probability density function of the unit circle. The mean parameter, *μ*, represents the mean of a given Von Mises, including one mean for positively signed reaches and one for negatively signed reaches (*μ*_1_ > 0, *μ*_2_ ≤ 0). Data were pooled across subjects in each RT bin and parameters were optimized separately in each bin using maximum likelihood estimation, minimizing the negative log likelihood with the MATLAB function *fmincon*. Fifty randomized starting parameter values were used for each fit to avoid local minima, and Aikake Information Criterion values (AICs) were either summed over all RT bins after the critical 7th bin or compared separately at each bin for model comparisons. Two models were compared: In the “Free *μ* model, both *μ* parameters could vary freely. In the “Fixed *μ* model, *μ* parameters were fixed at our *a priori* prediction of *μ_1_* = +90° and *μ_2_* = −90°. Thus, the Fixed *μ* model had two fewer degrees of freedom than the Free *μ* model.

## ACKNOWLEDGEMENTS

We thank Chandra Greenberg for help with data collection. We also thank Fulvio Domini, Justin Jungé, and Eugene Poh for helpful discussions and comments on the manuscript. This work was supported by the National Science Foundation (Graduate Research Fellowship to S. D. McDougle) and the National Institute of Neurological Disorders and Stroke (Grant R01 NS- 084948 to J. A. Taylor).

